# Distributions of host heterogeneity in susceptibility show signatures of pathogen geographic structure in an insect baculovirus

**DOI:** 10.64898/2026.06.18.732481

**Authors:** Arietta E Fleming-Davies, Savannah S Shields, Jacob Fletcher, Wilnelia Recart, David J Paez

## Abstract

Segregated variation between populations is a fundamental evolutionary process leading to parasite specialization, yet the resulting impacts on infection heterogeneity within populations are theoretically and empirically understudied. We asked whether the distribution of host susceptibility to infection within populations carries the signatures of geographic structure from pathogen local adaptation, maladaptation, or generalism in a nuclear polyhedrosis virus that infects the Gulf Fritillary butterfly *Dione vanillae*. For this virus, there is genetic support for two geographically distinct groups within San Diego County, based on whole genome sequencing of 16 virus isolates. Reciprocal laboratory infections showed evidence of two contrasting viral life history strategies: a ‘generalist’ phenotype that consistently infected variable hosts and a ‘specialist’ that performed slightly better in its local host population. As predicted by our theoretical model, the more consistent infection displayed by the generalist across populations corresponded to lower heterogeneity in susceptibility within populations, modeled as gamma distribution. Furthermore, the generalist phenotype was collected over a wider geographic range despite having a tenfold-lower mean infection rate than the specialist, suggesting that a strategy of more consistent infection provides key fitness advantages across diverse host populations. Intriguingly, when there is variation in host susceptibility, interpretations of pathogen local adaptation are dose-dependent. Measuring infectivity across multiple doses enables estimation of the whole distribution of susceptibility, which provides more reliable identification of pathogen specialization to its local host. Our work demonstrates how trait distributions and not only their mean values can carry quantifiable signatures of eco-evolutionary processes in interspecific interactions.

## Introduction

Coevolutionary processes are crucial in host-pathogen interactions due to the rapid rate of evolution typical of these interactions and the strength of selection on both pathogens and hosts. Host-pathogen coevolution with limited gene flow across landscapes can lead to local adaptation, in which pathogen specialization results in higher infection rates in sympatric compared to allopatric host-pathogen interactions (Blanquart *et al*. 2013; Lively & Dybdahl 2000).

Conversely, pathogen local maladaptation occurs when *host* adaptation decreases susceptibility to local pathogens, resulting in lower infection rates for sympatric than for allopatric host-pathogen pairs (Kaltz *et al*. 1999). In contrast, a generalist strategy favors more consistent infection across genetically variable hosts, typically at the cost of lower infection in any particular host type or species (Antonovics *et al*. 2013). Previous work has long recognized the importance of genetic variation within populations in pathogen local adaptation (Kawecki & Ebert 2004; Ridenhour & Nuismer 2007). Yet this work is largely separate from the substantial disease ecology literature on continuous variation in host susceptibility and its consequences for disease dynamics (but see Best *et al*. 2011; Lion *et al*. 2022). Host heterogeneity has been well-described by models that include continuous distributions of susceptibility within a host population, typically as Gamma or Beta distributions (Ben-Ami *et al*. 2010; Dwyer *et al*. 1997; Gomes *et al*. 2022; Hass *et al*. 2014; Langwig *et al*. 2017). Here, we examine how quantitative variation in infection changes across geographically structured populations, and find that the signatures of pathogen local adaptation, maladaptation and or generalism can be captured from changes of trait distributions in these commonly-used disease ecology models.

While gene-for-gene coevolution is common in many host-pathogen systems (Agrawal & Lively 2002), continuous or quantitative variation in transmission-related traits such as infectiousness (e.g. superspreaders; Lloyd-Smith *et al*. 2005) and host susceptibility to infection are also ubiquitous (Barrett *et al*. 2009). Here we focus on the distribution of susceptibility to infection, which can be estimated from laboratory dose response experiments (Ben-Ami *et al*. 2010). Different pathogens exhibit different susceptibility distributions when tested in the same host population, and thus changes in the heterogeneity of susceptibility result from selection on both pathogens and hosts (Fleming-Davies *et al*. 2015). In population models, increased heterogeneity in susceptibility can strongly reduce the size of epidemics, even in populations with identical mean transmission rates (Gomes *et al*. 2022; Hawley *et al*. 2024; Páez & Fleming-Davies 2020; Rose *et al*. 2021; Tuschhoff & Kennedy 2024). Within-species continuous variation in susceptibility is widespread across host-pathogen interactions, from invertebrate viruses (Dwyer *et al*. 1997) to bacterial diseases of wild birds (Hawley *et al*. 2024) to human pathogens (Gomes *et al*. 2022). Such heterogeneity arises from both genetic and environmental factors, including diet and prior exposure (Ben-Ami *et al*. 2010; Elderd *et al*. 2008; Hawley *et al*. 2024; Langwig *et al*. 2017; Páez *et al*. 2017). Despite the ubiquity of heterogeneity in susceptibility and the recognized importance of variation in coevolutionary processes, there is a lack of empirical and theoretical work describing how susceptibility distributions change across geographically structured host and pathogen populations.

Theory shows that local adaptation is favored by low gene flow in both interacting species, high pathogen virulence, and constant selection over time (Bono *et al*. 2017; Gandon & Michalakis 2002; Kawecki & Ebert 2004; Morgan *et al*. 2005). In empirical work, however, the strongest predictor of pathogen local adaptation is higher rates of pathogen gene flow compared to hosts, with other theorized factors, such as host taxonomy, relative generation time, and virulence, having little predictive power (Greischar & Koskella 2007; Hoeksema & Forde 2008). While this discrepancy in findings might be due to a lack of statistical power, attempts to identify factors in study design have struggled to find consistent general patterns (Hoeksema & Forde 2008), although some recommendations have emerged, such as conducting multiple reciprocal transplants and using mixed effects models (Kawecki & Ebert 2004). Here we argue that this mismatch between theoretical predictions and empirical results could also be due to the overwhelming focus on average trait differences instead of comparing the entire trait distribution between populations. Indeed, populations may have similar average phenotypes and yet be locally adapted if reciprocal transplant experiments show fitness differences (Kawecki & Ebert 2004). Thus, it is unsurprising that the signatures of local adaptation have been more reliably captured by comparing patterns of genetic variance between populations, for example by determining whether the magnitude of additive genetic variation in a quantitative traits among population is different from neutral expectations (e.g. comparing Q_st_ and F_st_) (Antoniazza *et al*. 2010; Leinonen *et al*. 2013; Savolainen *et al*. 2007; Whitlock & Guillaume 2009). We propose a method to reconcile this discrepancy between genetic and phenotypic results by quantifying distributions of infection susceptibility within populations using multiple infection doses. This technique is similar to methods using additive genetic variation in quantitative traits (Leinonen *et al*. 2013; Whitlock & Guillaume 2009), but without requiring explicit genetic information that is difficult to obtain in some systems.

We used a theoretical model to predict how distributions of susceptibility to infection might change under pathogen local adaptation, maladaptation, or generalism. We then applied this framework to estimate empirical distributions of infection within and across natural populations of an insect-baculovirus interaction. The nucleopolyhedrosis virus (NPV) that infects the larvae of the Gulf Fritillary butterfly *Dione vanillae* (Ribeiro *et al*. 2019; Rodríguez *et al*. 2011), is similar to many other Lepidoptera NPVs that exhibit high genetic and phenotypic diversity within host populations (Fleming-Davies *et al*. 2015; Hodgson *et al*. 2001). Our field collections found strong geographic structure in viral genomes at a small spatial scale, clustering isolates into two strains with different life history strategies. The ‘generalist’ phenotype was more consistent at infecting variable hosts both within and across populations, but at the cost of lower mean infectivity in all host populations. In contrast, the specialist strain had higher mean infection rates, particularly in its sympatric host population, but more variable infection rates both within and across populations. Importantly, the signature of pathogen local adaptation or generalism was more reliably captured in the continuous distributions of susceptibility to infection, as described by the heterogeneity terms, than by the differences in mean infection rates alone.

## Model

We modeled heterogeneity of infection as a gamma distribution of susceptibility to infection, susceptibilities *x* as follows (Ben-Ami *et al*. 2010; Langwig *et al*. 2017):

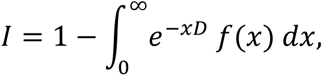

where *I* is the proportion of infected hosts, *D* is dose, and *f(x)* is the gamma probability distribution function. Assuming constant pathogen exposure over a fixed time interval, the proportion of infected hosts at dose *D* can be written in terms of the mean infection rate 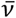 and coefficient of variation *C* (Ben-Ami *et al*. 2010):

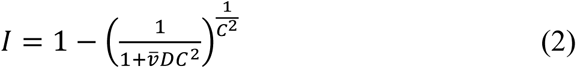

Both mean 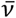 and heterogeneity of infection C^2^ are pathogen traits subject to selection, with higher mean or lower heterogeneity values leading to more infected hosts during an outbreak (Gomes *et al*. 2022; Hawley *et al*. 2024; Langwig *et al*. 2017) and thus higher pathogen fitness (Páez & Fleming-Davies 2020). We extended this theory to predict effects of pathogen local adaptation, maladaptation, and generalism on the distribution of susceptibility to infection (Fig 1). Specifically, a locally adapted pathogen in its sympatric host (Fig 1 top left panel) should have high mean and low variation in susceptibility because of directional selection on the pathogen to specialize on that host (Bulmer 1976). In allopatric populations, where local selection has not acted on the foreign pathogen, we expect a lower mean and higher variation in susceptibility. Alternatively, pathogen maladaptation results from directional selection acting on the host, resulting in both a low mean and a narrower distribution of susceptibilities (lower *C^2^*) in the sympatric association. In contrast, the ability of a ‘generalist’ pathogen strain to more consistently infect variable hosts should be evident in both lower heterogeneity *C^2^* within populations and as more similar infection between allopatric and sympatric populations. Thus, we expect the same intermediate mean and low heterogeneity *C^2^* across all populations (Figure 1, far right panels).

**Figure 1:**
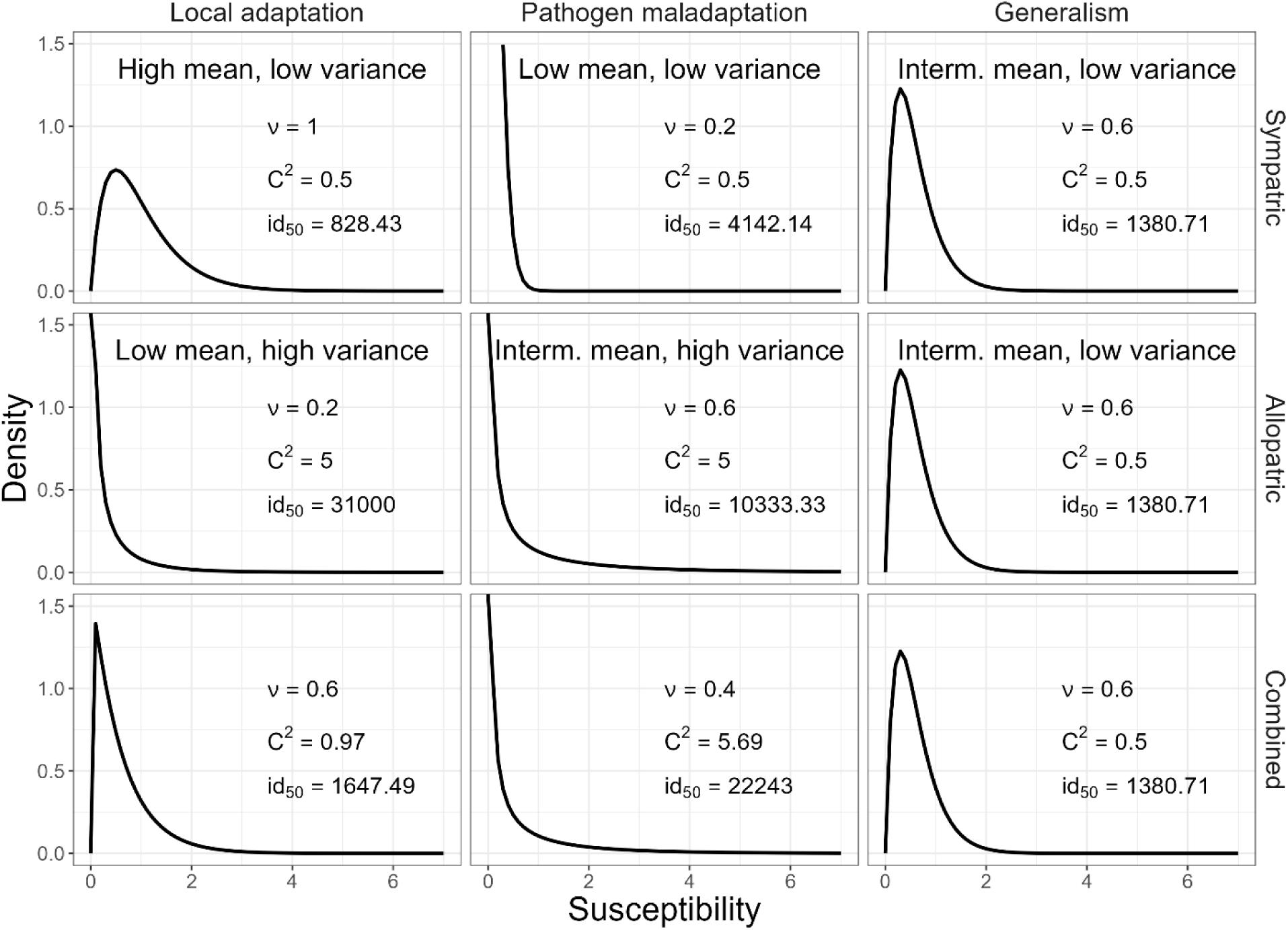
Hypothetical gamma distributions of susceptibility in sympatric, allopatric, and combined populations (rows top to bottom) for pathogen local adaptation, maladaptation, and generalism (columns left to right). Values of mean susceptibility 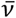 and heterogeneity of susceptibility C^2^ were set by assumptions for scenarios in the top two rows. Gamma distributions were computed for the combined population (bottom row) by averaging the sympatric and allopatric gamma distributions using a moment matching approach.

Using the above predictions for local adaptation, maladaptation, and generalism, we next considered how the above parameters change in an average or combined population consisting of both subpopulations (allopatric and sympatric) equally represented. We used a moment matching method to combine the allopatric and sympatric Gamma distributions, using the Welch-Satterthwaite approximation (Stewart *et al*. 2007) that allows the sum of *n* Gamma distributions with different shape and scale parameters *k_i_* and *θ_i_* to be described by a Gamma (distribution with shape 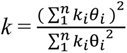 and scale 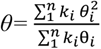. In the two population case, we multiply by the scalar ½ before adding the distributions to average across the allopatric and sympatric populations, giving a new Gamma distribution of susceptibility with a mean infection 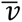 that is simply the average of the two single population means, and a combined heterogeneity term 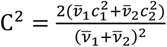.

Importantly, the signature of geographic structure (either local adaptation or maladaptation) is visible in the combined population as a higher heterogeneity value C^2^ when compared to the generalist pathogen (Figure 1). Furthermore, while the locally adapted strain in its sympatric population has the highest fitness of the three pathogen strategies (indicated by the lowest LD50 value, the dose to infect 50% of the population for a lethal pathogen), the generalist strain has the highest fitness in the combined population, consistent with a generalist strategy.

## Methods

### Collections of NPV isolates

To quantify viral genetic variation across San Diego County, we collected host *Dione vanillae* larvae from cultivated *Passiflora spp.* plants (Figure 1). Field-collected larvae were raised individually on surface-disinfected *Passiflora spp.* leaves until death from virus or pupation. Virus-killed insects were stored at −20°C for viral occlusion body isolation using a standard protocol (See SI methods).

### Viral genome sequencing

After viral DNA extraction following standard methods for NPVs (see SI), library preparation and whole genome *de novo* sequencing were performed at a commercial facility (Center for Aquaculture Technologies, San Diego, CA, USA). Specifically, constructed DNA libraries were subjected to 150bp paired-end sequencing on an Illumina HiSeq platform. See SI methods for quality control details. *De novo* assembly was performed using the metaviralSPAdes pipeline in SPAdes version 3.15.5 (Bankevich *et al*. 2012). Viral related sequences were identified using VirSorter2 version 2.2.4. Genome assembly resulted in 1-11 contigs per sample (size range: 18132 to 123013bp). BLAST searching of the NCBI database suggested matches with high coverage and percent identity to an Alphabaculovirus, *Dione juno* nucleopolyhedrovirus isolate Araguari-MG (Ribeiro *et al*. 2019). In each of our samples, the largest assembled scaffold had a large percent identity to the Alphabaculovirus genome and was therefore used in subsequent analyses. We performed a multiple sequence alignment of these scaffolds (n=16 isolates) to the reference Alphabaculovirus genome in MAFFT with default parameters (Katoh *et al*. 2002), and generated neighbor-joining consensus trees using maximum likelihood methods (Jukes-Cantor model) in IQTree (Nguyen *et al*. 2014). Following whole genome analyses, we examined geographic patterns of genetic variation among isolates in six key viral genes (See Table S2 for gene functions and sizes).

### Dose response experiments

We conducted reciprocal laboratory dose response experiments using larvae and virus isolates from two populations representative of the major geographic grouping of viral genomes within San Diego County: North County (collected from Encinitas, CA); and the City of San Diego (collected from central San Diego; Figure 1). Healthy 4^th^ instar larvae used in infections were second generation lab-bred offspring of butterflies from each population. See SI methods for insect rearing and collection details. Larvae from each population (North County hosts: n=12 larvae per dose; San Diego City hosts; n=18 larvae per dose) were exposed to four pathogen doses (a water control or 100, 1000, or 10,000 occlusion bodies) of one of two virus isolates: BFF-004 from North County or GRM-060 from central San Diego City (Figure 1). Lab infections were conducted using a modified droplet feeding method with 3 uL occlusion bodies in water on 8mm diameter leaf disks (Elderd *et al*. 2008).

### Model fitting and statistical analysis

We used a Bayesian approach to estimate the parameters 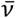 and C that describe the Gamma distribution of host susceptibility (eqs. 1-2) from the laboratory infection data. All models were fit in R, using MCMC with a Gibbs sampler (Casella & George 2016), assuming a binomial error distribution (infected/uninfected) and non-informative lognormal priors. We used Bayesian Information Criteria (BIC) to compare models in which each host and pathogen combination had a unique mean infection rate 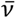 and heterogeneity of infection *C* to simpler models in which pathogen strains varied but host populations were not considered or vice versa, and to a null model with no host or pathogen differences and no heterogeneity (Table S2). We also used a general linear model assuming binomially distributed data to test for effects of pathogen and host population on infection in the equivalent homogeneous model (i.e., no variation of susceptibility within a host population; glm function in R).

We summarized the overall pathogen performance across doses in the heterogeneity model as the dose at which 50% of individuals are infected (LD_50_) from the dose response curves for each host *×* pathogen combination. LD_50_ was computed as 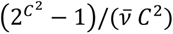, derived from eq. 2 by setting I=0.5 and solving for dose *D* (Ben-Ami *et al*. 2010). We then used the ID_50_ values to test for local adaptation. We followed previous work to compute the strength of local adaptation *E* in sympatric *s* and allopatric *a* host populations, *E*=ln(*X*_s_/*X*_a_) (Hoeksema & Forde 2008). Thus, *E* is a unitless effect size that takes positive values for pathogen local adaptation (i.e., higher performance in sympatric hosts) and negative values for maladaptation (i.e., higher performance in allopatric hosts). We take the inverse to account for the fact that lower values of ID_50_ indicate higher pathogen fitness, giving *E*=ln (ID50_a_/ID50_s_). To determine if either of the pathogen strains specialized on their local host, we computed *E* for each pathogen strain and across strains.

## Results

### Geographic variation in virus isolates

Analysis of viral genomes revealed extensive genetic variation within San Diego County (Figure 1). Pairwise identity between isolates ranged from 94.3% to 99.8% over the whole 122,383 bp genome (Table S1). Phylogenetic analyses revealed that virus genotypes grouped into two major clades which strongly corresponded with geographic structure (Figure 1). We will refer to these groups as the “City of San Diego strain” and the “North County strain” due to the high degree of genetic difference between these clades and the evidence of phenotypic differences described below. Within the North County collection sites (BFF and OLE; Figure 1B), we observed only the North County strain. While the City of San Diego strain was only collected in the central sites (CDO, ARM, GRM; Figure 1B), the North County strain was present at all sites but one (ARM, with the lowest sampling effort).

Sequence analyses of key viral life history genes controlling within-host processes and environmental persistence were also consistent with this geographic structure (Figure 2); most of the non-synonymous substitutions and insertion-deletions in these genes occurred between the two main clades (Table S1). However, we observed additional variation in several genes within the City of San Diego strain (Table S1, Figure 2), which contributed to substantial standing variation within collection sites. For example, isolates collected at the GRM site included four variants (non-synonymous predicted amino acid sequences) of the chitinase gene *v-chi* (Fig 2B; Table S1). In general, City of San Diego collection sites had higher standing variation present, including genotypes from both clades (upper right heatmap quadrants, Figure 2). In contrast, we found no non-synonymous polymorphisms among isolates collected at the North County sites for any of the genes examined, despite a similar collection effort (Table S1).

**Figure 2:**
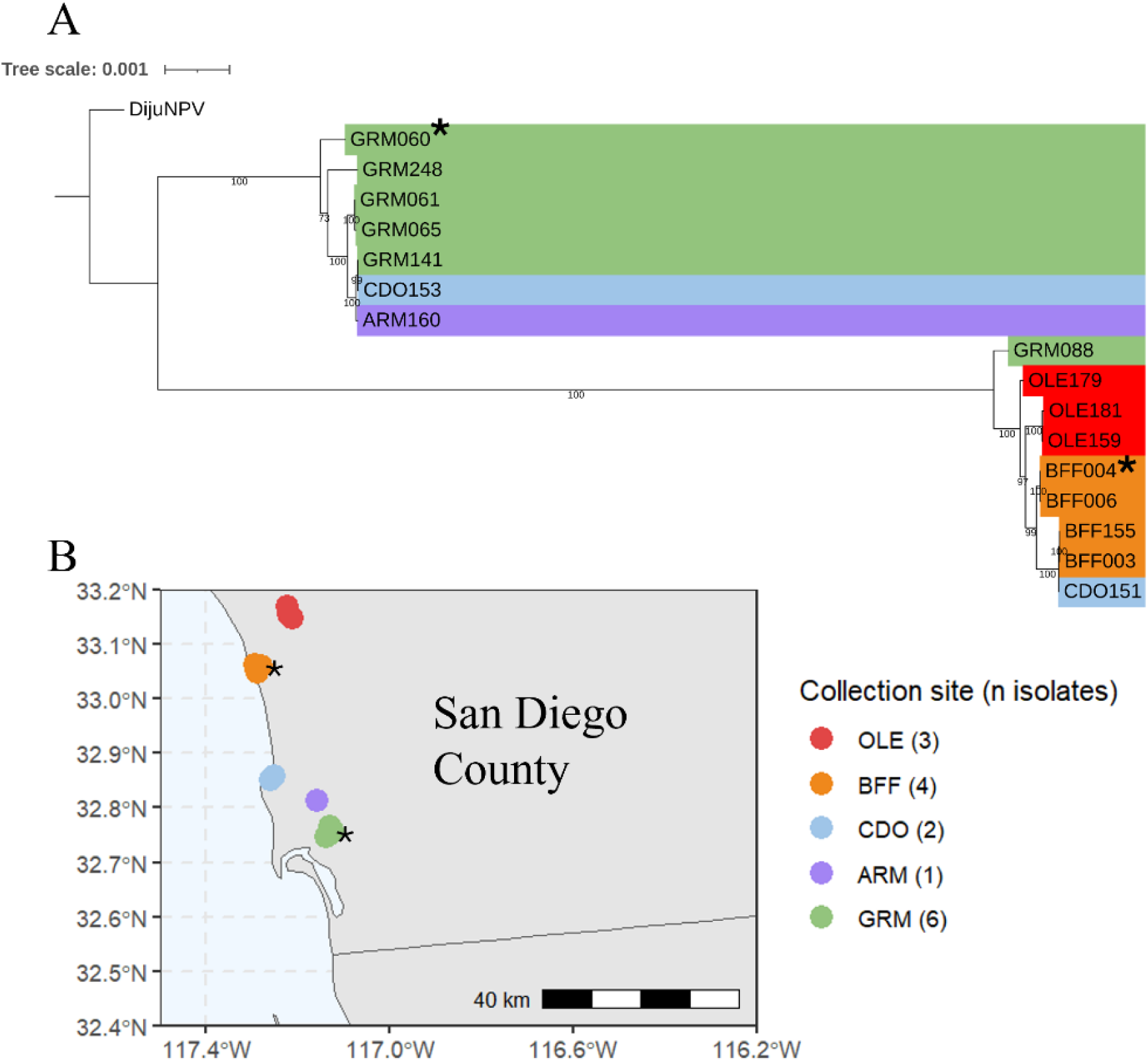
Virus isolates collected in San Diego County clustered in two distinct genetic groups: a “City of San Diego strain” (top clade in A) found only in the central region, and a “North County strain” (lower clade in A) that was the only strain collected in the North County sites (OLE, BFF; red and orange dots in B) but was also collected at several central sites (GRM, CDO). Note that similar collection efforts were made at the different sites and thus the variation in number of isolates per site reflect variation in plant and/or virus prevalence. Asterisks mark the isolates used in local adaptation dose response experiments. DijuNPV is the reference genome of the NPV collected in Dione juno (NC_076692) (Ribeiro *et al*. 2019). Phylogenetic tree was constructed using multiple sequence alignment of whole genomes; numbers below branches give maximum likelihood support values for tree structure.

### Phenotypic differences between virus strains

Virus strains representative of the North County and San Diego City clades differed in their dose response curves and the resulting distributions of susceptibility (Figure 3, strain by dose interaction, LR= 4.47, df=1, p=0.03, GLM assuming binomial distribution, Table S2). To test whether the distributions of host susceptibility match the theoretical expectations of local adaptation, maladaptation or generalism, we compared susceptibility distributions for each strain between sympatric, allopatric, and combined (i.e., ignoring host population) conditions. While the San Diego City strain (GRM-060) had the highest mean and lowest variation of susceptibility in its sympatric host population, as predicted for a specialist strategy of adaptation to its local host, the North County strain (BFF-004) demonstrated a phenotype more consistent with generalism, producing similar distributions of susceptibility across all populations. As predicted by our theoretical model for a locally adapted or specialist strain (Fig 1, leftmost column), the San Diego City strain exhibited higher mean susceptibility and smaller variation in susceptibility (Fig. 4) when infecting its sympatric host population compared to its allopatric population (median and 66% HPD credible intervals: 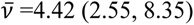 and *C^2^*=4.95 (3.42, 4.99) in sympatric; 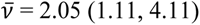; *C^2^*= 4.12 (3.91, 6.33) in allopatric), although differences between host populations were small when compared to differences between virus strains (Table S3). The log ratio of LD_50_ in allopatric and sympatric populations for the San Diego City strain was also consistent with local adaptation of that strain to its sympatric host (Fig. 5; E = 1.21 (0.997, 2.36); estimate and 95% CI). In contrast, the North County strain had similar or worse infectivity in its sympatric host population compared to its allopatric population (E = −0.36 (−1.53, 0.06); 95% CI overlaps 0). Overall, the North County strain matched predictions of our theoretical model for a generalist strategy, with more consistent infection of variable hosts both within populations (as measured by lower heterogeneity of susceptibility, the mean-scaled coefficient of variation) and between populations (as measured by more similar values of mean and CV in different host populations). When dose response data were pooled across host populations (i.e. combined case in Fig. 1), the San Diego City virus strain (GRM-060) demonstrated almost a ten-fold higher mean infection rate compared to the North County strain (San Diego: median and 66% credible intervals: 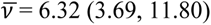; North County: 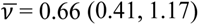), but also slightly more variable infection, as represented by the mean-scaled squared coefficient of variation (San Diego City C^2^= 5.19 (4.43, 6.07); North County C^2^ = 4.11 (3.29, 5.21)). Speed of kill was very similar across strains (Figure S1).

**Figure 3:**
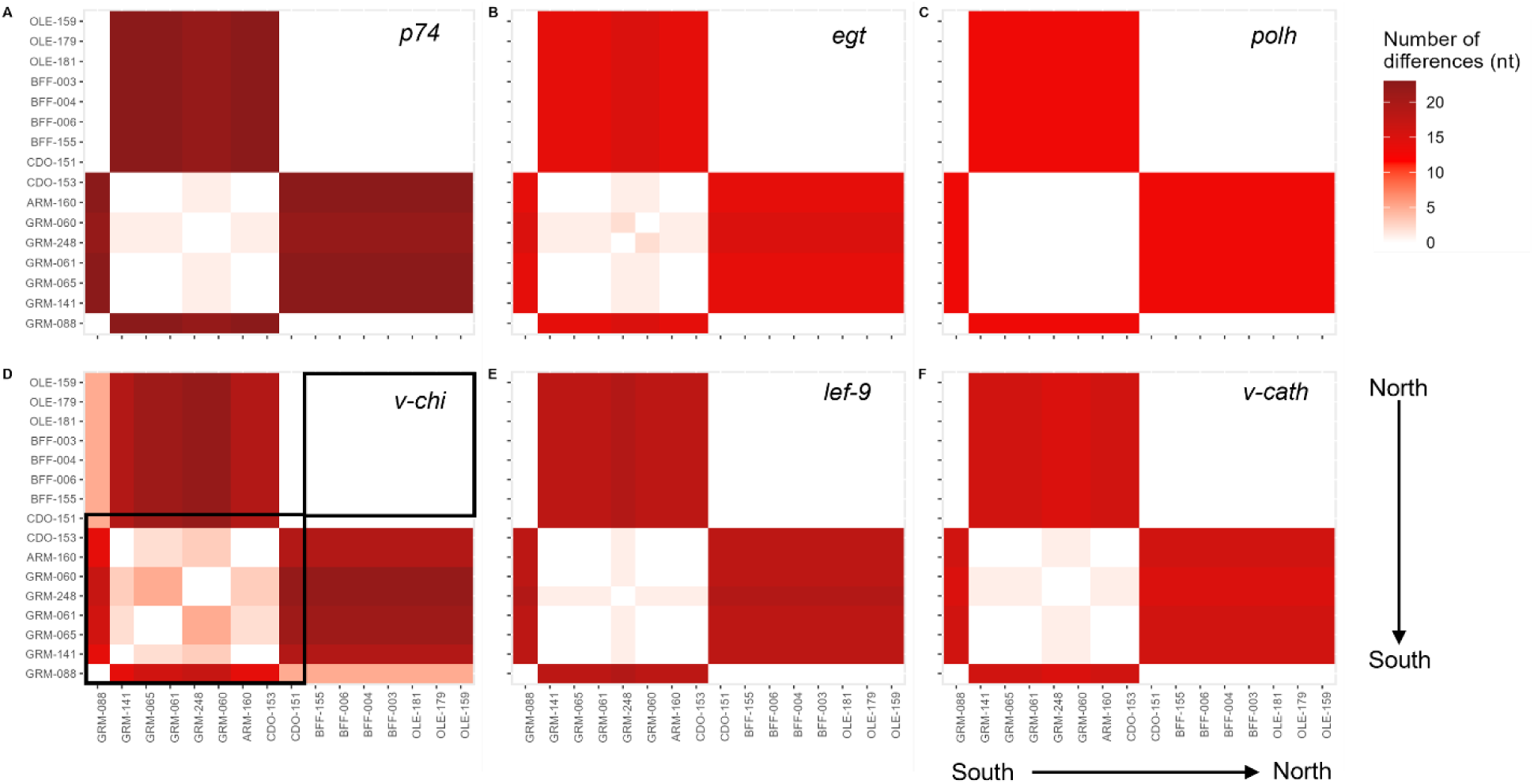
Heat maps of genetic distance (number of nucleotide differences) for pairwise comparisons between field-collected isolates for key viral genes. See Table S1 for gene sizes and functions. Sites (three letter code) in each heat map are ordered top to bottom and right to left by geographic distance from the northernmost site (OLE; Figure 1b). Isolates within a collection site are ordered arbitrarily. White indicates an identical DNA sequence. All six genes displayed strong genetic structure between the North County and City of San Diego clades, and more genetic variation within City of San Diego sites (lower left quadrant; marked by square in D) compared to North County sites (upper right quadrant).

**Figure 4:**
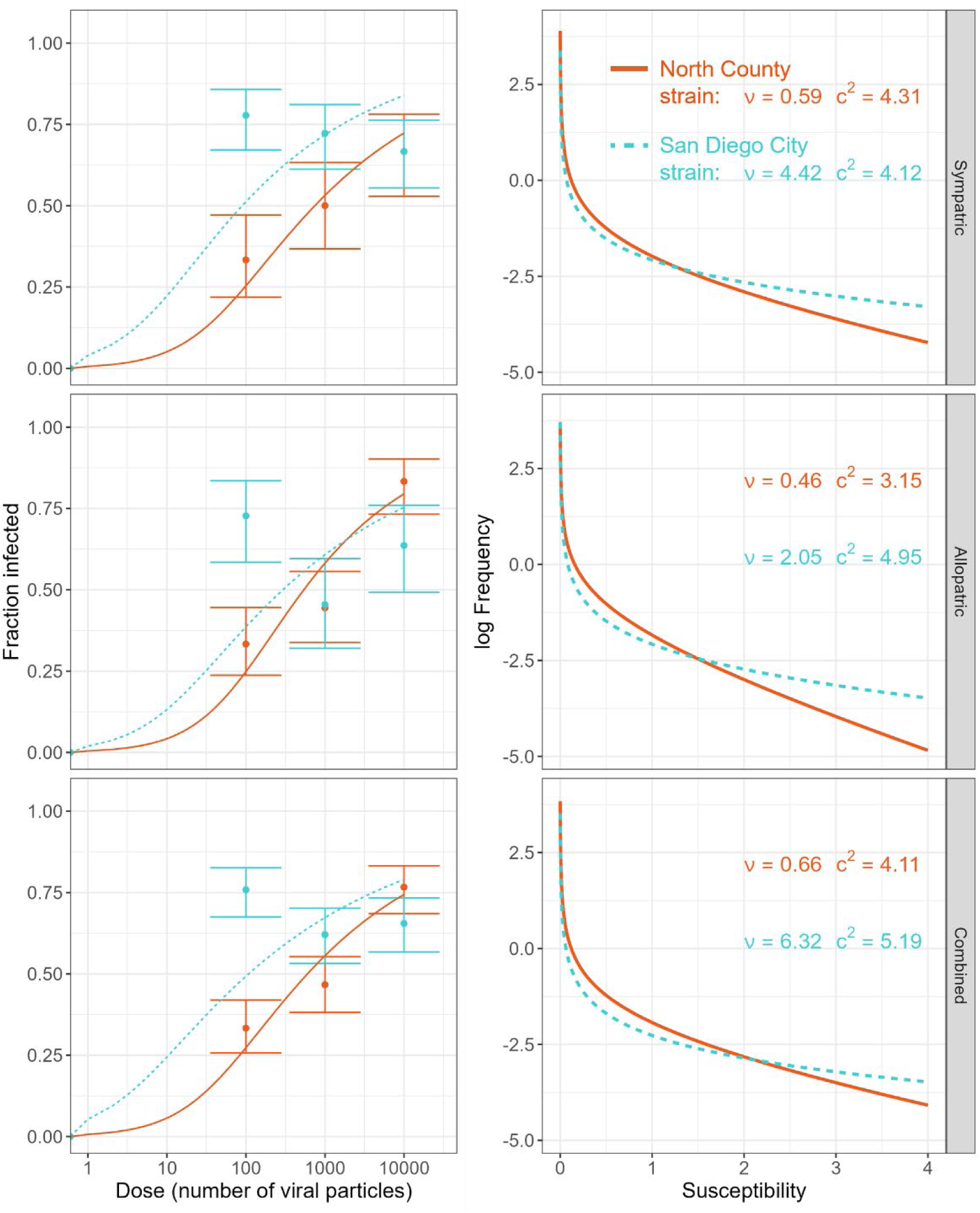
Dose response data (left panels) and fitted Gamma distributions of susceptibility (right panels) for virus strains from North County (solid orange lines) and San Diego City (dotted green lines) clades in sympatric, allopatric, or combined host populations (rows).

**Figure 5:**
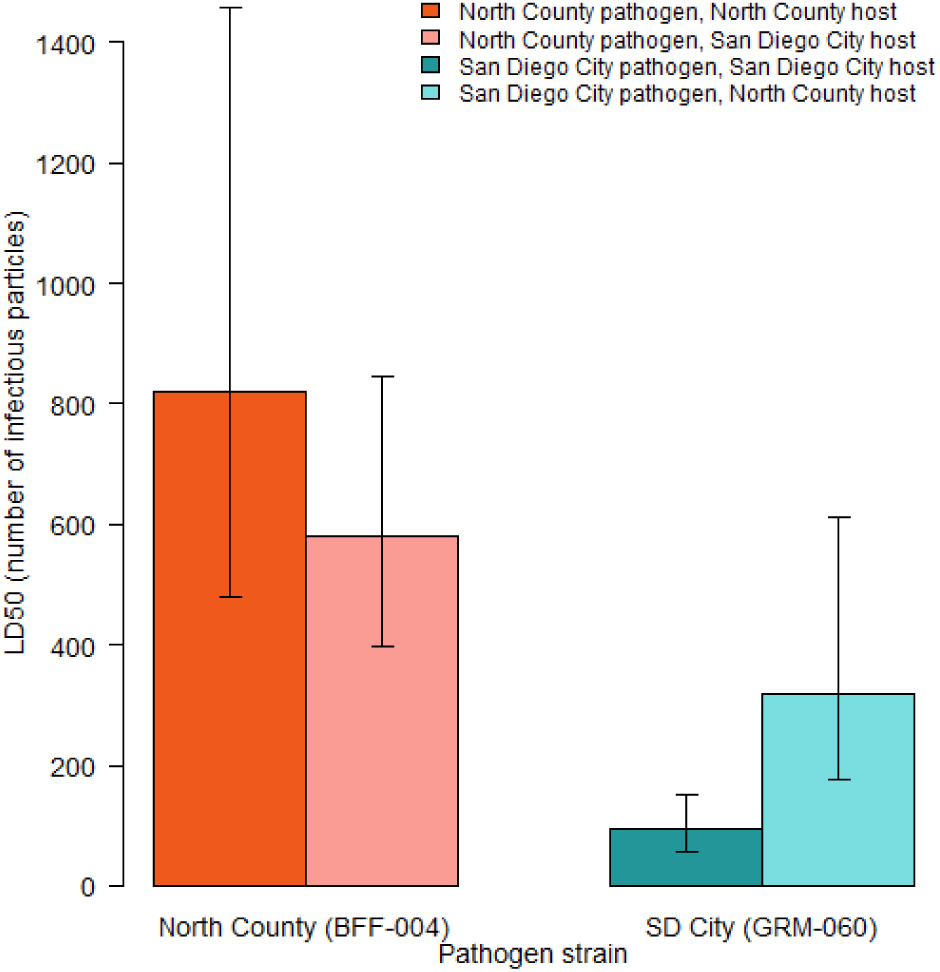
The San Diego City pathogen strain had higher infection, indicated by a lower dose required to infect 50% of individuals (LD_50_), compared to the North County pathogen strain in either sympatric (darker bars) or allopatric (lighter bars) host populations. Bars and error bars give medians and 66% HPD credible intervals (equivalent to 1SE) from posterior distributions of parameters from the model fitting Gamma distributions of susceptibility.

In contrast to the strong differences between viral strains, we found weak to no effects of host population on infection (Figures 3 & 4, host population effect LR=0.928, df=1, p=0.33, GLM assuming binomial distribution, Table S3). There was also no evidence of a host by pathogen population interaction in the raw infection data as predicted if local adaptation of each pathogen strain to its host population was maintaining this geographic structure (LR=0.259, df=1, p=0.61; GLM with binomial data; Table S3).

## Discussion

Characterizing susceptibility distributions provides a powerful approach to quantify specialization or generalism in infectious diseases that is more consistent with the evolutionary ecology of continuous traits. Local adaptation to discrete host types in host-pathogen interactions is often placed as one extreme in the continuum of pathogen specialization (Antonovics *et al*. 2013), but consistent infection *within* host types, as measured by reduced infection heterogeneity, is equivalent or an even more extreme case of specialization. Thus, quantifying distributions of susceptibility to infection provides a useful measure of pathogen specialization, particularly if the within-population results are consistent with the between-population results as we saw in this system. Under this framework, we found that the signatures of pathogen generalism or specialization to a local host were evident in the changing distributions of susceptibility within populations in the *D. vanillae*-NPV system. Specifically, the generalist North County pathogen strain was more consistent in its infection both within and across host populations; this strain had lower within-population heterogeneity of infection as well as more similar parameters across the two host populations (Figures 3B, 4). In contrast, the more specialized SD City strain was less able to infect a variable host population, as indicated by larger heterogeneity value C^2^ (Figure 3), and performed slightly better in its local SD City host due to both increased mean infection and decreased variation in that population, as predicted by our theoretical model if it were evolving to specialize on its local host (Fig. 1).

It is possible that the ubiquity of studies failing to find local adaptation (e.g, 54% of studies reviewed by Greischar & Koskella 2007) in part arises from continuous host heterogeneity in those systems, limiting the power to quantify the effects of adaptation on fitness-related traits using a single dose. Specifically, continuous variation in susceptibility introduces nonlinearities in the dose response curve, causing the relative infectivity values to vary with dose beyond the expected changes in variance of a binomial variable. For example, with a four-fold difference in mean susceptibility between sympatric and allopatric interactions 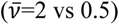 and heterogeneous susceptibility (C^2^=4 vs 2), infectivity at the lowest dose would support local adaptation (log ratio E=0.51) whereas the highest dose would indicate no local adaptation or even maladaptation (E=-0.03). In contrast, using LD_50_ from the heterogeneity model estimated with multiple doses would correctly support local adaptation (E= 0.98). This dependence on dose strengthens as heterogeneity increases; if all hosts are identical in their susceptibilities (C^2^=0), very high doses might not be informative, but the estimated local adaptation does not change sign. Furthermore, with continuous variation in susceptibility within populations, local adaptation can still occur under equal average infectivity between populations. For example, local adaptation can lead to smaller heterogeneity in susceptibility in the sympatric compared to the allopatric population, resulting in more consistent infection of hosts in that population and thus higher pathogen fitness. Dose response experiments allow us to quantify the heterogeneity signatures of selection without specific genetic information. Furthermore, allowing for host heterogeneity in disease ecology studies is a conservative assumption, given that it is widespread even in organisms with simple immune systems such as invertebrates (Ben-Ami *et al*. 2010; Dwyer *et al*. 1997). Recent theoretical work has explored new methods of approximating changes in the variance of trait distributions in scenarios including local adaptation, using an “oligomorphic approximation” to connect quantitative genetics and adaptive dynamics (Lion *et al*. 2022). To our knowledge, this exciting work has not yet been extended to empirical examples.

Our theoretical framework captures an ecological moment in time, and the distributions of these traits will then go on to affect the further coevolution of host and pathogen populations. In local adaptation theory, phenotypic and additive genetic variation allows for a fast response to selection, which can lead to rapid local adaptation and geographical differentiation (Falconer & Mackay 1983). Furthermore, in nature, heterogeneity C is influenced by environmental traits such as prior exposure as well as by genetic variation, thus adding another level of complication when predicting the response to selection. While the simplifying assumptions of our model ignore the additional complexity of coevolutionary change over time, the signature of these processes should still be present in the continuous distributions of susceptibility, making this simple approach a powerful first step at detecting the signatures of pathogen specialization to host populations.

Our results highlight how life-history tradeoffs of specialist and generalist infection strategies might promote two-strain pathogen coexistence. Specifically, the lower heterogeneity value of a more generalist strain provides an advantage to the otherwise disadvantageous lower average infection, promoting coexistence in a single host population (Fleming-Davies *et al*. 2015). The San Diego City strain had a mean transmission rate almost ten times higher than the North County strain, and yet the North County strain is found at low densities across a broader geographic range than the SD City strain. Importantly, our results suggest that definitions of generalist and specialist strategies should include measures of the distribution of susceptibility within and between the different host types (i.e., genotypes or species), in order to capture the consistency of infection among hosts. Most NPVs are quite specialized to one or several host Lepidoptera species (Cory & Myers 2003). The closest match to our *de novo* sequenced viral genomes (~96-98% identity; Table S1) was *DijuNPV*, the NPV that infects *Dione juno* larvae that share larval food plant species with *D. vanillae* in their native range (Ribeiro *et al*. 2019). Our results suggest that our collected isolates are the same virus species as *DijuNPV* and that there is high standing genetic variation at very small spatial scales, as our San Diego County strains were each more similar to the *DijuNPV* reference genome than they were to each other. Interestingly, we observed variable genome sizes among isolates as well as insertions in repetitive regions (Ribeiro *et al*. 2019) between North County and Central San Diego virus strains. This could be evidence of a ‘genomic accordion,’ first proposed in poxviruses, in which duplication of repetitive regions generates genetic diversity, then selection reduces genome size while retaining advantageous mutations in the expanded regions (Elde *et al*. 2012). Variation in genome size has been observed in other insect baculoviruses (Erlandson 2009) alongside evidence of positive selection (Simón *et al*. 2011), suggesting that this mechanism for rapid evolution of double-stranded DNA viruses might also occur in NPVs.

The strong spatial structure we observed in virus genotypes might also be due to geographic barriers to pathogen gene flow. Only the non-alate larval stage of the Gulf Fritillary butterfly is infected, limiting the range of virus dispersal. Mechanisms of long-distance dispersal of other NPVs are variable and understudied. In some NPVs, there is evidence of vertical transmission after sublethal infection (Ilyinykh 2019), which would allow volant adults to move pathogen genotypes over longer distances. In other NPVs, long-distance dispersal of the pathogen is thought to be primarily driven by ballooning of neonatal larvae and movement of larval populations as they defoliate large tracts of forest (Dwyer & Elkinton 1995). Avian and invertebrate predators of Lepidoptera also contribute to the spread of NPV in some systems (e.g. Entwistle *et al*. 1993; Lee & Fuxa 2000; Reilly & Hajek 2012), although *D. vanillae* adults are avoided by predators in North America (Ross *et al*. 2001). Finally, geographic structure in this urban wildlife virus might arise from human movement and even socioeconomic factors (Schell *et al*. 2020). If human movement of virus-contaminated plants for propagation is a method of dispersal, then human neighborhood structure could be reflected in pathogen geographic structure. More research is needed in the *D. vanillae*-NPV system to determine the mechanisms and geographic scale of pathogen dispersal. Local adaptation has been observed at the continental scale in other NPVs (Escribano *et al*. 1999; Simón *et al*. 2011). It is possible that this NPV is locally adapted at larger geographic scales such as across host subspecies (Halsch *et al*. 2020).

Quantifying local adaptation, generalism, and maladaptation from the distributions of susceptibility to infection has the key benefit of incorporating host and pathogen traits involved in their ecological interaction (McPeek 2017). Increased heterogeneity of susceptibility can lead to dramatic reductions in total number of individuals infected (Gomes *et al*. 2022; Hawley *et al*. 2024; Langwig *et al*. 2017; Páez & Fleming-Davies 2020) and have strong impacts on pathogen fitness (Fleming-Davies *et al*. 2015). Our understanding of local adaptation and specialization in infectious diseases would be improved by estimating distributions of susceptibility rather than relying on means. Experimentally, this is a simple change to multiple doses for each host *x* pathogen combination in ‘reciprocal transplant’ experiments across populations. Furthermore, multi-dose experiments to fit distributions of susceptibility within and across populations provide another powerful method to apply “distributional thinking” (Wetzel *et al*. 2023) to understand eco-evolutionary processes and detect the signatures of evolution in complex ecological systems.

## Acknowledgements

We thank former undergraduate students R Bernhardt, R Solis, and M Velesrubio for assistance with field collections, and M Martinez, M Lee, and T Winthrop for assistance in the lab. We also thank V Hohman and G Morse of USD and J Stannard and M Allen of The Center for Aquaculture Technologies for their advice and assistance with DNA extractions and genome sequencing. Funding for this work was provided by: NSF DEB CAREER #2145704 to PI Fleming-Davies; USDA-NIFA AFRI grant #2017-67015-26956 to PI Naish which partly supported D Páez’s work on this study; University of San Diego (USD) Office of Undergraduate Research; USD Department of Biology; a USD Faculty Research Grant; and the USD Honors College.

Any use of trade, firm, or product names is for descriptive purposes only and does not imply endorsement by the U.S. Government.

