## Supplemental Information for "Distributions of host heterogeneity in susceptibility show signatures of pathogen geographic structure in an insect baculovirus"

### Supplemental Methods:

*Study system:*

The Gulf fritillary butterfly, *Dione vanillae* (Nymphalidae; subgenus *Agraulis*) (Zhang et al. 2019; Penz 2022) is an introduced species at the study site in San Diego, CA, USA, where it has been established for several hundred years (Forister et al. 2018). Its distribution is primarily determined by the presence of the *Passiflora spp.* plants consumed by its herbivorous larvae (Halsch et al. 2020). In the urban environment of San Diego, larval food plants consist of cultivated ornamental and fruiting species such as *Passiflora caerulea*, *P. incarnata*, *P. edulis*, and various hybrids of the above (Copp and Davenport 1978, Passiflora Society International 2011). The host-specific nuclear polyhedrosis virus (AgvaNPV) that infects *D. vanillae* is a lethal baculovirus transmitted when larvae consume contaminated leaf material (Rodríguez et al. 2011). After death, the infected caterpillar's cuticle dissolves, dripping virus onto leaves that are then consumed by conspecifics, thus spreading the infection. The virus is contained in large protein-coated occlusion bodies that protect it from the environment and allow for viral persistence outside the host (Miller 1997). Occlusion bodies are visible at 400X magnification, allowing for easy detection of infection and field collection of isolates.

*Field sites and collection of virus isolates*

Field sites on private land were identified by community members or through the iNaturalist app. During each site visit, all host plants and surrounding areas were searched for larvae for approximately 30 person-minutes. Any virus-killed larvae observed in the field were collected on site, and living larvae were brought back to the lab and raised individually on surface-disinfected *Passiflora spp.* leaves until death from virus or pupation (see *Insect rearing* below). After death from confirmed NPV infection in the lab, virus-killed insects were collected in 1.5mL tubes and stored at -20°C.

*Insect collection and rearing for laboratory experiments*

Larvae used in dose response infection experiments were the lab-raised offspring of field collected individuals. The parental generation was collected as healthy larvae (n=10) from free-living populations at each site (BFF in North County and GRM in the City of San Diego; Figure 1). Field collected larvae were raised individually on surface-disinfected *Passiflora spp* leaves until pupation. Presence of the NPV inhibits host molting hormones and thus successful pupation indicates that no active virus is present (Miller 1997). Pupae were then transferred to single population mating colonies. After emergence, adult butterflies were allowed to mate freely within their population. Adults and larvae were raised in a growth chamber in an NPV-free clean space at 25°C and 70% humidity on a 12-hour light/dark cycle, on surface-disinfected leaf material from *Passiflora* "Quasar", a hybrid of *P. caerulea* 'Constance' *x P. pallens* (Passiflora Society International 2011). To control for plant effects on infection, all leaf material for rearing and infections was a randomized mix from five greenhouse-grown plants, (Shikano et al. 2017).

To control for developmental effects on infection (Cory and Myers 2003), all larvae were exposed to pathogens at the 4th instar.

*Isolation of Viral Occlusion Bodies*

NPVs are unique in that when death of the host larva occurs, the individual is typically >95% viral DNA by weight (Miller 1997). This allows for very simple collection and extraction of virus isolates. Viral occlusion bodies were isolated from virus-killed insects in water following a standard protocol (Kennedy and Dwyer 2018). For each field-collected NPV-killed larva, up to 400µl of water was added to the sample to bring the total volume up to 1000 µl. The 1.5 mL tube was then vortexed for two minutes, and 200µl of this solution was transferred to a fresh epitube, and the remaining solution was stored in the freezer at -20°C. 800µl of DI water was added to the 200µl occlusion body solution, homogenized by vortexing for 1-2 minutes. The solution was centrifuged at 4,000 RPM for 3 minutes to obtain a pellet of occlusion bodies. The supernatant was removed, and the occlusion body pellet was resuspended in 800µl of DI water again, followed by vortexing for 1-2 minutes and centrifuged at 4,000 RPM for 3 minutes. The resulting pellet containing occlusion bodies was suspended in 500 µl DI water, then dissolved by vortexing for 1-2 minutes and stored at -20°C for use in laboratory infections or DNA extraction for whole genome sequencing.

*Viral DNA Extraction*

To extract viral DNA, occlusion body pellets were resuspended in 1mL 1M $Na2CO3$ to simulate the alkaline environment of the insect gut (after Rodríguez et al. 2011a). After vortexing, 20µL Proteinase K was added to each sample to ensure protein digestion. Samples were incubated overnight (16 hours) at 37°C and viral DNA was extracted from 300 µl of the incubated occlusion using the Qiagen Blood and Tissue DNAeasy Kit from Step 1d onwards. The quality of extracted DNA was assessed using concentration and A260/280 values measured using a nanodrop. Extracted DNA was then stored at -80°C to maintain quality. Polymerase chain reaction (PCR) was used to verify the presence of AgvaNPV DNA in each virus sample by amplifying the *p-74* gene, which is essential for the baculovirus life cycle and thus highly conserved among NPV species (Faulkner et al. 1997). We used the primers and cycle used to amplify the P-74 gene described by Rodriguez et al. (2011): p74-Reverse: GAAACTCGCGCGGAAACAT, p-74 Forward: CGCGGGTGCGANAGCATG. The PCR cycle was 94 °C for 2 min (1cycle); 92°C for 10s, 48°C for 12s, 72°C for 1min 30s (20cycles); 92°C for 10s, 60°C. PCR amplicons of 661/664 bp were run on a 1% agarose gel. Samples confirmed by PCR and gel electrophoresis were sent for sequencing.

*Bioinformatics Analysis*

In addition to the phylogenetic analysis using whole viral genomes outlined in the main text, we also quantified differences in gene sequences and predicted amino acid sequences for six viral genes crucial in the viral life cycle (see Table S2 for list of genes and functions). Individual genes were identified in Geneious (www.geneious.com). The first step was to map all isolate consensus sequences (longest scaffold) to the reference *Dione juno* Alphabaculovirus genome (Ribeiro et al. 2019), which was ~95-98% identical to our samples. The second step was to extract the individual genes identified in that reference genome. DNA sequences for each isolate were aligned using MAFFT to generate matrices of pairwise differences for each gene (heatmaps in Figure 3). Next, the predicted amino acid sequences for each of the six genes, were generated in Geneious using the 'translate' tool for all potential open reading frames. To estimate the number of non-synonymous substitutions, insertion-deletions, and amino acid variants for the six genes among the 16 field-collected virus isolates we aligned the amino acid sequence from the most likely open reading frame to reference proteins using Geneious. This was repeated for each of the six viral genes (Table S2).

*Speed of kill estimates*

Infection outcome and day of death was noted by visual inspection (i.e. melanized, 'melted' phenotype of virus-killed larva), and confirmed with necropsies to check for presence of occlusion bodies under a light microscope at 400X (Miller 1997). Previous work has documented non-linear tradeoffs between the probability of infection and the speed of host kill (Fleming-Davies et al. 2015; Páez et al. 2017). Therefore, we also measured the speed of host kill for the reciprocal dose response infection experiment by recording the day of larvae death, defined as the time at which no larval movement was observed after gentle stimulation, correlated with increased melanization and swelling of the cuticle.

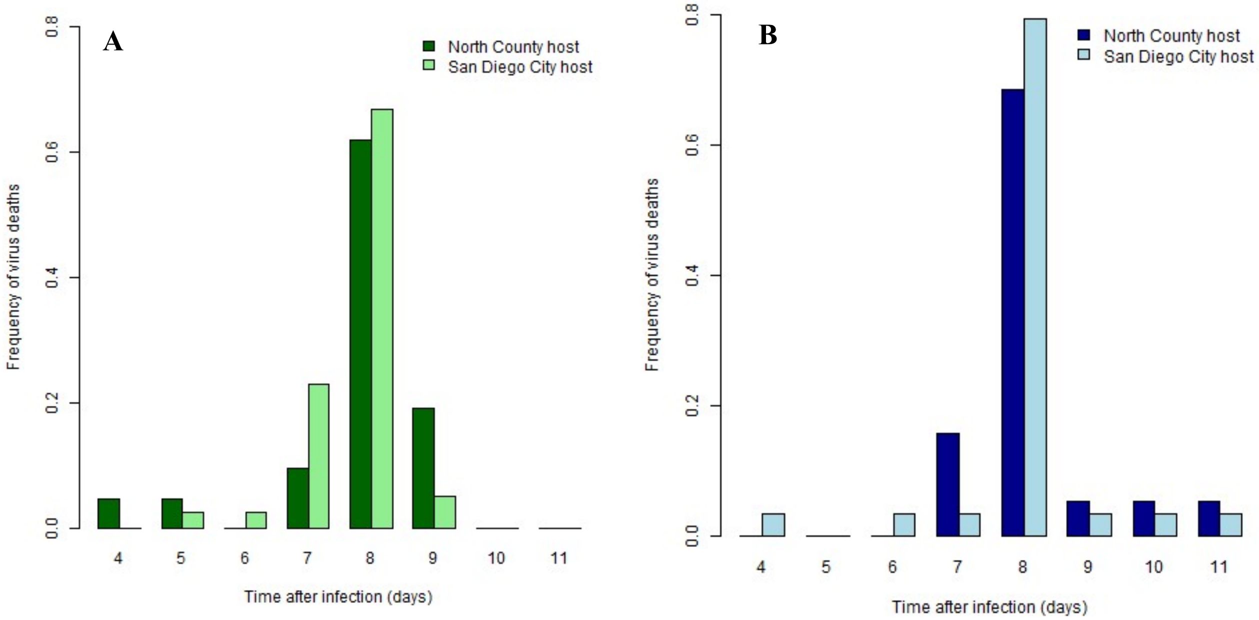


**Figure S1:** Speed of kill of San Diego City (A; GRM-060) and North County (B; BFF-004) virus strains in laboratory infections of North County (dark bars) and San Diego City (light bars) host populations. Relative frequencies are number of larvae dying on that day divided by the total number of virus-killed larvae for each virus strain and host combination (San Diego City strain: n=59; North County strain: n=47) to allow comparisons between host and pathogen populations despite unequal sample sizes. The higher-virulence City of San Diego strain killed hosts slightly but not significantly faster (mean ± SE: 7.72 ± 0.12 days from infection to death) than the lower-virulence North County strain (8.03 ± 0.15 days; ANOVA; effect of strain: $F_{1,101}=3.65$, $p=0.06$). There was also a non-significant trend towards faster speed of kill at higher doses (ANOVA; effect of dose: $-2.8 \times 10^{-5}$; $F_{1,101}=1.06$, $p=0.30$) as expected from the literature (e.g. Hodgson et al. 2001), and no effect of host population (ANOVA; $F_{1,101}=0.12$ $p=0.74$).

**Table S1:** Pairwise percent identity between all collected NPV isolates and the reference genome (DijuNPV; NC076692). Multiple sequence alignment using MAFFT, default settings. Isolates used in experimental infections are highlighted.

| | Diju NPV | OLE-159 | OLE-179 | OLE-181 | BFF-003 | BFF-004 | BFF-006 | BFF-155 | CDO-151 | CDO-153 | ARM-160 | GRM-060 | GRM-248 | GRM-061 | GRM-065 | GRM-141 | GRM-088 |
|---|---|---|---|---|---|---|---|---|---|---|---|---|---|---|---|---|---|
| **DijuNPV** | | 95.42 | 95.86 | 95.59 | 96.10 | 96.16 | 96.16 | 96.10 | 96.00 | 98.44 | 97.88 | 97.74 | 97.89 | 98.11 | 98.06 | 98.44 | 96.53 |
| **OLE-159** | 95.42 | | 97.96 | 99.63 | 97.72 | 97.85 | 97.85 | 97.73 | 97.73 | 94.82 | 94.39 | 94.78 | 94.79 | 95.03 | 94.79 | 94.82 | 98.29 |
| **OLE-179** | 95.86 | 97.96 | | 98.22 | 98.13 | 98.17 | 98.17 | 98.13 | 98.13 | 95.30 | 94.77 | 95.17 | 95.17 | 95.51 | 95.26 | 95.30 | 98.74 |
| **OLE-181** | 95.59 | 99.63 | 98.22 | | 97.99 | 98.12 | 98.12 | 97.99 | 97.99 | 95.07 | 94.55 | 95.03 | 95.04 | 95.29 | 95.04 | 95.08 | 98.56 |
| **BFF-003** | 96.10 | 97.72 | 98.13 | 97.99 | | 99.22 | 99.22 | 99.69 | 99.47 | 95.60 | 95.07 | 95.42 | 95.32 | 95.49 | 95.40 | 95.60 | 99.26 |
| **BFF-004** | 96.16 | 97.85 | 98.17 | 98.12 | 99.22 | | 99.79 | 99.32 | 99.01 | 95.66 | 95.13 | 95.55 | 95.38 | 95.62 | 95.53 | 95.66 | 99.30 |
| **BFF-006** | 96.16 | 97.85 | 98.17 | 98.12 | 99.22 | 99.79 | | 99.32 | 99.00 | 95.66 | 95.13 | 95.55 | 95.38 | 95.62 | 95.53 | 95.66 | 99.30 |
| **BFF-155** | 96.10 | 97.73 | 98.13 | 97.99 | 99.69 | 99.32 | 99.32 | | 99.47 | 95.60 | 95.07 | 95.42 | 95.32 | 95.49 | 95.40 | 95.60 | 99.26 |
| **CDO-151** | 96.00 | 97.73 | 98.13 | 97.99 | 99.47 | 99.01 | 99.00 | 99.47 | | 95.50 | 94.97 | 95.32 | 95.22 | 95.39 | 95.30 | 95.50 | 99.16 |
| **CDO-153** | 98.44 | 94.82 | 95.30 | 95.07 | 95.60 | 95.66 | 95.66 | 95.60 | 95.50 | | 99.12 | 98.65 | 99.02 | 99.44 | 99.39 | 99.79 | 96.06 |
| **ARM-160** | 97.88 | 94.39 | 94.77 | 94.55 | 95.07 | 95.13 | 95.13 | 95.07 | 94.97 | 99.12 | | 98.10 | 98.47 | 98.88 | 98.83 | 99.12 | 95.53 |
| **GRM-060** | 97.74 | 94.78 | 95.17 | 95.03 | 95.42 | 95.55 | 95.55 | 95.42 | 95.32 | 98.65 | 98.10 | | 98.34 | 98.67 | 98.63 | 98.66 | 95.95 |
| **GRM-248** | 97.89 | 94.79 | 95.17 | 95.04 | 95.32 | 95.38 | 95.38 | 95.32 | 95.22 | 99.02 | 98.47 | 98.34 | | 98.89 | 98.64 | 99.02 | 95.93 |
| **GRM-061** | 98.11 | 95.03 | 95.51 | 95.29 | 95.49 | 95.62 | 95.62 | 95.49 | 95.39 | 99.44 | 98.88 | 98.67 | 98.89 | | 99.74 | 99.44 | 96.18 |
| **GRM-065** | 98.06 | 94.79 | 95.26 | 95.04 | 95.40 | 95.53 | 95.53 | 95.40 | 95.30 | 99.39 | 98.83 | 98.63 | 98.64 | 99.74 | | 99.39 | 95.93 |
| **GRM-141** | 98.44 | 94.82 | 95.30 | 95.08 | 95.60 | 95.66 | 95.66 | 95.60 | 95.50 | 99.79 | 99.12 | 98.66 | 99.02 | 99.44 | 99.39 | | 96.06 |
| **GRM-088** | 96.53 | 98.29 | 98.74 | 98.56 | 99.26 | 99.30 | 99.30 | 99.26 | 99.16 | 96.06 | 95.53 | 95.95 | 95.93 | 96.18 | 95.93 | 96.06 | |

**Table S2:** Polymorphisms between and within North County and San Diego City viral genomes in key life history genes. Well-characterized genes were chosen to represent key processes at each stage of the infection cycle: infection of host gut cells after consumption (*p74*), control of host behavior and molting after infection (*egt*), host dissolution (*pquit, pcat*), and occlusion body production (*polh*). Only changes in predicted amino acids (ie, non-synonymous substitutions or insertion-deletions) are listed as polymorphisms; for total genetic variation in these genes see Fig 2. Residue number is given in parentheses. Number of variants is the number of different predicted amino acid sequences for that gene among all isolates.

| **Gene (length in nt)** | **Function** | **Polymorphisms present, SD version given first (residue number)** | **Number of variants** |
|---|---|---|---|
| *p74* (1,944 nt) | Envelope protein; permits entry to host gut cells in initial infection (Kuzio et al. 1989) | Lysine to glutamic acid (159), valine to isoleucine (531), glutamic acid to lysine (582), valine to methionine (604) | 2 (North County; City of San Diego) |
| *egt* (1,503 nt) | Ecdysteroid uridine 5′-diphosphate (UDP)–glucosyltransferase; inactivates host molting hormone; controls 'zombie' climbing behavior in infected host (Hoover et al. 2011) | Indel: two aspartic acid inserted in North County strain (194-5), glycine to serine in GRM060 (299), glycine to glutamic acid (428) | 3 (North County; GRM060; all others in City of San Diego clade) |
| *v-chi* (1,676 nt) | Chitinase; digests host chitin (Hawtin et al. 1997) | Leucine to arginine (51), Alanine to valine (129) in GRM060 and GRM248, lysine to asparagine (214), threonine to asparagine (277) in all North County clade except GRM088, lysine to asparagine (374), lysine to glutamic acid (475), asparagine to lysine (555) | 4 (GRM088; all others in North County clade; GRM060/GRM248; all others in City of San Diego clade) |
| *v-cath* (975 nt) | Cathepsin; digests host protein (Hawtin et al. 1997) | Lysine to arginine (34), glycine to alanine (95) | 2 (North County; City of San Diego) |
| *lef-9* (1482 nt) | Late expression factor; regulates genes for occlusion body production (Lu and Miller 1995) | Alanine to valine (9),proline to serine (238), alanine to aspartic acid (384), asparagine to aspartic acid (466) | 2 (North County; City of San Diego) |

| | | | |
|---|---|---|---|
| *polh* (738) | Polyhedrin; produces outer protein matrix in occlusion bodies (Van Iddekinge et al. 1983) | Tyrosine to histadine (17), leucine to phenylalanine (51), histadine to tyrosine (95), lysine to glutamine (106), glutamic acid to lysine (128), phenylalanine to leucine (133), phenylalanine to leucine (143), alanine to threonine (151), leucine to phenylalanine (163), leucine to phenylalanine (206), lysine to glutamic acid (209) | 2 (North County; City of San Diego) |

**Table S3:** Effect sizes and likelihood ratio tests from general linear model analysis of laboratory dose response data of infections with City of San Diego and North County pathogen and host populations. GLM assumed binomial data (infected/uninfected) with logit link. Asterisks denote significant effects at the α=0.05 level.

| Effect | Effect size | LR | df | p-value |
|---|---|---|---|---|
| Host population | SD city: 0.150 | 0.9277 | 1 | 0.33 |
| Pathogen population | SD city: 0.736 | 3.00 | 1 | 0.08 |
| **Dose*** | **$2.43 \times 10^{-4}$** | **26.6** | **1** | **<0.001** |
| **Pathogen population x dose*** | **SD city: $-1.46 \times 10^{-4}$** | **4.47** | **1** | **0.03** |
| Pathogen x host population | SD city x SD city: 0.25 | 0.2593 | 1 | 0.61 |
